# Differentiating *Peromyscus leucopus* bone marrow-derived macrophages for characterization of responses to *Borrelia burgdorferi* and lipopolysaccharide

**DOI:** 10.1101/2024.12.19.628444

**Authors:** Christopher C. Wells, Tanja Petnicki-Ocwieja, Shumin Tan, Stephen C. Bunnell, Sam R. Telford, Linden T. Hu, Jeffrey S. Bourgeois

## Abstract

Currently, most tools utilized in host-pathogen interaction studies depend on the use of human or mouse (*Mus musculus*) cells and tissues. While these species have led to countless breakthroughs in our understanding of infectious disease, there are undoubtably important biological processes that are missed by limiting studies to these two vertebrate species. For instance, it is well-established that a common North American deermouse, *Peromyscus leucopus*, has unique interactions with microbes, which likely shape its ability to serve as a critical reservoir for numerous zoonotic pathogens, including a Lyme disease spirochete, *Borrelia burgdorferi*. In this work, we expand the immunological toolkit to study *P. leucopus* biology by performing the first differentiation of deermouse bone marrow to macrophages using *P. leucopus* M-CSF producing HEK293T cells. We find that *P. leucopus* BMDMs generated through this method behave broadly very similarly to C57BL/6J macrophages generated with L-929 supernatant, though RNA sequencing revealed modest differences in transcriptomic responses to *B. burgdorferi* and lipopolysaccharide. In particular, differences in *Il10* induction and caspase expression were observed between the species.

## INTRODUCTION

*Peromyscus* deermice are the most abundant mammals in North America [1] and serve as a crucial reservoir for many microorganisms that cause human disease [2–8]. While microbes with pathogenic potential are ubiquitous in nature, numerous scholars have noted that the ability for “pathogens” to cause disease depends on the interplay between host and microbial processes, with insufficient or excessive inflammation proving detrimental to the host [9]. While this damage-response framework is frequently applied to human-microbe interactions, these concepts are also critical among wild vertebrates, as symptomatic disease can impair the ability to obtain food or evade predation. Thus, it is perhaps unsurprising that studies with *Peromyscus leucopus* have revealed an unusual ability for the rodent to limit inflammation and resist disease or death in response to numerous pathogens, including *Borrelia hermsii* [10, 11], Powassan Virus [2], and Sin Nombre Virus [12].

In particular, *Borrelia burgdorferi* sensu stricto (*B. burgdorferi*), the predominant Lyme disease spirochete in North America, is prototypical of this phenomenon [13–20]. Following infection with *B. burgdorferi*, *P. leucopus* do not exhibit any major symptoms associated with human Lyme disease, including swelling of major joints or febrile responses [17–20], though limited carditis is observed [21]. Despite this limited inflammation, *P. leucopus* and related deermice are actually better at restricting *B. burgdorferi* burden and rodent-to-tick transmission of the spirochete compared to laboratory mice (*Mus musculus*) [21–23], including strains that display Lyme disease-like illness. Thus, *P. leucopus* appears able to restrict *B. burgdorferi* without maladaptive inflammation through currently unknown mechanisms.

While *B. burgdorferi* lack lipopolysaccharide (LPS) [24], the interactions between *P. leucopus* and LPS have also been subject to substantial study [10, 11]. When attempting to induce septic death from *E. coli* LPS injection, researchers noted *P. leucopus* require a dose many-fold higher than that of *M. musculus* to achieve lethality [11]. Additionally, when comparing *P. leucopus* and *M. musculus* under LPS stimulation, discriminant analysis of relative gene expression in whole blood demonstrates that *P. leucopus* utilizes *Il1b* responses to LPS stimulation in comparison to *Ifng* responses observed in *M. musculus* [10]. Similar to their interactions with *B. burgdorferi*, the immunological mechanisms behind *P. leucopus*’ pro-survival immune response to LPS are poorly understood, though a role for macrophages has been proposed [10].

The study of *P. leucopus* transcriptomic responses to *B. burgdorferi* and LPS have largely been performed using tissues from infected animals [10, 11, 19, 21, 25] or skin fibroblasts [11, 26, 27]. While powerful in providing initial descriptive data regarding the responses to bacterial stimuli, these methods have some limitations. Bulk RNA sequencing studies produced from tissue harvested following infection have highlighted pathways important to host immune responses, but lack the ability to attribute those changes to a specific cell population. Furthermore, the heterogeneity of these samples may bias the findings from sequencing towards bystander populations rather than the less abundant immune cell population, potentially producing a type-two error in the analysis of the results. More specific inquiries into skin-derived fibroblasts have the benefit of greater sample homogeneity compared to whole-tissue and are related to skin-based pathogenicity and *Borrelia* spp. transmission; however, they may not be representative of the central immune traits of the species because of their nature as a tissue specific mesenchymal cell population. Notably, there have been very few described methods to generate a specific immune cell population for *ex vivo* study of host-pathogen interactions with *P. leucopus*—though one study was able to isolate and stimulate peripheral macrophages following pretreatment of rodents with Freund’s adjuvant [28]. Here, we describe the development of *P. leucopus* macrophage colony-stimulating factor (M-CSF) secreting cells for the purpose of differentiating macrophages from flushed bone marrow. We characterize the macrophages and find that at the transcript level, they behave very similarly to *M. musculus* (C57BL/6J) bone marrow-derived macrophages (BMDMs) differentiated using supernatant from L-929 cells, both during basal conditions and during stimulation with either *B. burgdorferi* (strain B31) or LPS (derived from *Salmonella enterica* serovar Minnesota R595), though some differences were observed in *Il10* expression and caspase expression. Together, these studies shed light on *P. leucopus* interactions with Gram-negative microbes as well as lay the groundwork for future studies attempting to understand *P. leucopus*-pathogen interactions.

## METHODS

### Rodent Maintenance, Use, and Ethical Statement

All animal procedures were approved by the Tufts University-Tufts Medical Center Institutional Animal Care and Use Committee (IACUC, Protocol Numbers B2021-84, B2024-50). Euthanasia was performed in accordance with guidelines provided by the American Veterinary Medical Association (AVMA) and was approved by the Tufts University IACUC. Rodents were maintained by Tufts University Comparative Medicine Services. C57BL/6J mice purchased from Jackson Laboratories (#000664). *P. leucopus* were provided by Dr. Sam Telford and are derived from northeastern and midwestern captured rodents. The *P. leucopus* colony has been closed since 1994, maintained in microisolator cages, and is specific-pathogen free, confirmed by regular sentinel testing. All rodents used in this study were 7-14 weeks old, with both species sex-matched and approximately age-matched within each experiment.

### Experimental Design, Statistics, and Data Exclusion

Two male and two female rodents from each species were used for BMDM generation. For each experiment, one rodent from each species (age and sex matched) was utilized for BMDM generation and used for all experiments in this manuscript. Thus, all data represent four independent experiments, with one exception. One male *P. leucopus* and C57BL/6J was excluded from the RNA sequencing as multiple samples from that experiment had low RNA yields and quality, likely resulting from a technical issue during RNA isolation. Statistical tests performed throughout the paper are described in the appropriate figure legend. All statistics were performed using either GraphPad Prism 10 software or R version 4.3.1.

### Bacterial Strains and Culture

*Borrelia burgdorferi* strain B31 (Hu Lab AD strain, a non-clonal strain routinely reisolated from sequential passages through C3H/HeN mice to maintain virulence) was utilized for all stimulations and microscopy. Spirochetes were cryopreserved in 25% glycerol and stored at -80°C. The spirochete was grown in Barbour-Stonner-Kelly II (BSK-II) complete media [29] at 37°C until it reached 1 x 10^7^-5 x 10^7^ spirochetes/mL, as quantified by darkfield microscopy using a Petroff-Hauser Chamber. In all experiments a multiplicity of infection of 100 was used to infect cells. Plasmids (**Table 1**) were maintained in DH5α (Thermo Fisher) *E. coli. E. coli* were grown in LB media with 50 μg/mL kanamycin (Sigma K1377-5G) or 100 µg/µL ampicillin (Fisher BP1760-5).

**Table 1:** Plasmids used in this study.

| **Table 1: Plasmids used in this study** |  |  |  |  |
| --- | --- | --- | --- | --- |
| Laboratory ID | Construct Name | Description | Antibiotic Resistance | Source |
| JSB_Ec09 | SB_5266 | pLK4 Cloning Vector | Kanamycin | [30] |
| JSB_Ec11 | JSB_Ec11_Csf1 | *P. leucopus* Csf1 cassette cloned into JSB_Ec09_SB_5266 | Kanamycin | This Work, Plasmid map in **Supplemental File 1** |
| JSB_Ec12 | pMD2_G_addg | VSV-G Envelope | Ampicillin | AddGene Plasmid 12259 |
| JSB_Ec13 | psPAX4_Min | Gal-Pol-RRE | Ampicillin | This Work |
| JSB_Ec14 | psPRTT_REV | Rev-T2A-Tat | Ampicillin | This Work |

### Cell Lines, Plasmid Constructs, and Transfection/Transduction

L-929 cells (ATCC CCL-1) were cultured in DMEM + 10% heat inactivated FBS. HEK293T from ATCC (CRL-3216) were used to generate *P. leucopus* M-CSF producing cells. HEK293T cells were grown in DMEM with 4.5 g/L glucose, L-glutamine, sodium pyruvate (Corning MT10013CV) supplemented with 10% heat inactivated FBS (R&D Systems 1304537A). A lentivirus construct for expression of *P. leucopus*-M-CSF (JSB_Ec11_Csf1, **Table 1**) was transfected alongside three additional constructs (JSB_Ec12_pMD2_G_addg, JSB_Ec13_psPAX4_Min, JSB_Ec14_psPRTT_REV, **Table 1**) to generate lentivirus. JSB_Ec11_Csf1 was generated by cloning into into pLK4. pLK4 CMV mYFP MCS IRES-Puro (SB_5266) is a modular variant derived from the pLEX-MCS (Open Biosystems, Grand Island, NY) in the Bunnell lab [30]. VSV-G pseudotyping vector pMD2.G (AddGene Plasmid 12259) developed by Didier Trono. The lentiviral packaging vector psPAX4 (SB_5804) was derived from psPAX2 (AddGene Plasmid 12260) by deleting the sequences encoding REV and TAT. The accessory vector psPRTT (SB_5805) was assembled by installing a synthetic gBlock (IDT) encoding REV-T2A-TAT into the psPAX2 backbone. Together psPAX4 and psPRTT constitute a split backbone 4th gen packaging system developed in the Bunnell Lab. Two days post-transfection, viral supernatant was centrifuged at 200xg for five minutes, filtered through a 0.45 μm PDVF filter, and used to transduce HEK293T cells. This cell pool was then grown in 1 µg/mL puromycin selection to purify a M-CSF producing population. M-CSF-producing cells were maintained, cryopreserved, and used as uncloned pools following transduction.

### Bone Marrow Isolation and Differentiation

Rodents were euthanized with CO_2_ and C-spine dislocation following Tufts University IACUC guidelines Bone marrow was flushed from the femur with sterile DMEM as previously described [31]

To generate *M. musculus* differentiation medium, L-929 cells were cultured for two passages. During the third passage the L-929 culture was allowed to reach >80% confluency, and two days after achieving 80% density the cell culture supernatant was collected and frozen at - 20°C for later use as conditioned media for murine BMDM differentiation. Murine progenitors were cultured on 100 x15 mm^2^ plastic Petri dishes for 5-7 days in DMEM media supplemented with 30% L-929 cell-conditioned medium and 20% FBS. Alternative differentiation was carried out with 20 ng/mL recombinant murine M-CSF (Peprotech, #315-02) in DMEM + 20% FBS.

To generate *P. leucopus* M-CSF, transduced HEK293T cells were passaged to approximately 50% confluence in the presence of 1 µg/mL puromycin and allowed to reach 80% confluency. The culture was continued for two additional days and supernatant was collected, filtered through a 0.22 µm PES syringe filter, and stored at -20°C for use in BMDM differentiation. Bone marrow from *P. leucopus* was differentiated in DMEM supplemented with 30% HEK293T^M-CSF^ cell supernatant and 20% heat inactivated FBS in 100 x 15 mm^2^ sterile dishes or in 6-well plates.

For both species, 1 x 10^6^ cells were plated in 100 x 15 mm dishes with 10 mL of differentiation media, and 1 x 10^6^ cells in 12 mL differentiation media in 6-well plates with 2 mL per well. Cells were supplemented with an additional 0.5 volumes of their original plating at days two days and four of differentiation. On day six of differentiation, cells were lifted via cell-scraper and transferred to DMEM + 10% FBS. During differentiation, phase contrast microscopy images were captured on days zero, three, and six of differentiation in order to track cell morphology changes.

### Flow Cytometry

Throughout differentiation, cells were scraped and pelleted and washed in FACs buffer (PBS, 2% FBS, 2 mM EDTA). Cells were resuspended in 1 mL FACs buffer and stained with 1 µL LIVE/DEAD Fixable Aqua (ThermoFisher L34965) for 30 minutes in the dark at room temperature. Cells were washed in FACs buffer, fixed with 4% formaldehyde for 30 minutes, washed, and stored at 4°C before analysis on a Cytek Aurora flow cytometer.

### RT-qPCR

Throughout differentiation, RNA was collected by scraping cells into media, pelleting cells at 300xg, and resuspending in 700 µL of Qiazol (Qiagen 79306). RNA was isolated using the miRNAeasy isolation kit (Qiagen 217084). cDNA was synthesized from isolated RNA using the ImProm-II Reverse Transcriptase kit (Promega A3801). qPCR was completed using the iTaq Universal SYBR Green Supermix (Biorad 1725124). Reactions were performed using a Biorad CFX Connect Real-Time System. Cycling parameters were one cycle of 95°C for 15 minutes, 40 cycles of 95°C for 30 seconds, 60°C for 30 seconds, and 72°C for 30 seconds, and one cycle of 95°C for 1 minute, 55°C for 1 minute. Genes of interest were normalized to expression of *18s* in *M*. *musculus* and *P. leucopus* samples (**Table 2**).

**Table 2:**
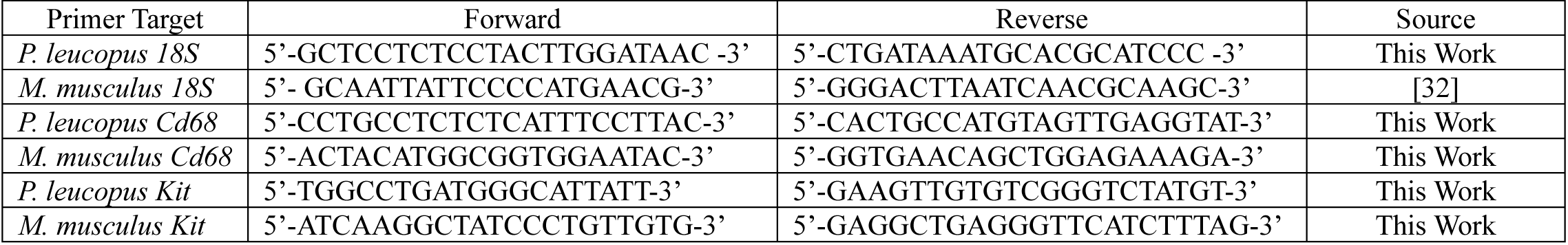
Oligonucleotides used in this study.

### *B. burgdorferi* Phagocytosis Assay and Confocal Microscopy

Round coverslips were sterilized in 70% ethanol and placed in the wells of a 24-well plate. Differentiated and lifted macrophages were plated onto coverslips at 3.0 x 10^5^ per well (6.0 x 10^5^ cells/mL). Cells adhered overnight and were then infected with *B. burgdorferi* B31, MOI 100 resuspended in 300 µL of complete DMEM for 5, 15, or 30 minutes. Coverslips were fixed in 2% paraformaldehyde in PBS at room temperature for 1 hour. Fixed cells were washed and permeabilized in blocking buffer (PBS with 2% goat serum and 0.1% saponin) overnight at 4°C. Cells were stained overnight at 4°C with mouse anti-*B. burgdorferi*-FITC 1:500 (Abcam ab69219). followed by rabbit anti-LAMP1 1:100 (Novus NB120-19294) for 2 hours at room temperature and a secondary. anti-rabbit Alexa Fluor 647 – far red (ThermoFisher A21244) for 2 hours. Coverslips were washed mounted with Prolong Gold DAPI Mountant

Images were captured with a Nikon A1R confocal microscope with a 63X oil objective. All images were captured while zoomed in 4X from the original field using the NIS Elements Software from Nikon. Captured images were evaluated quantitatively by blinded counters measuring the number of macrophage unassociated *B. burgdorferi*, coiled and phagocytosed *B. burgdorferi*, and LAMP1 colocalized *B. burgdorferi* as previously described [33–36].

### Phagosomal Maturation

In order to measure phagosome acidification, 5-(and-6)-carboxyfluorescein beads were generated according to previous protocols [37]. Carboxyfluorescein fluorescence is pH-sensitive when excited at 490 nm but not at 450 nm (emission at 520 nm in both cases), enabling ratiometric measurement of pH changes[37–39]. In order to measure phagosome proteolysis, DQ-BSA/Alexa Fluor (AF) 594 beads were generated according to previous protocols [38]. Proteolytic processes that occur as the phagosome matures hydrolyze the protein to single, dye-labeled peptides, relieving the quenching of the BODIPY dye. AF594 fluorescence is constitutive and unaffected by phagosomal maturation and allows ratiometric measurement of DQ-BSA fluorescence changes[37–39].

Carboxyfluorescein and DQ-BSA assays were carried out following prior protocols [39]. Differentiated, lifted macrophages were plated in black-walled TC-treated polystyrene 96 well plates (Corning 353376) 2.0 x 10^5^ cells per well and allowed to adhere overnight at 37°C, 5% CO_2_. Cells were washed 3 times using cuvette buffer (PBS, 5% FBS, 5 mM dextrose, 1 mM CaCl_2_, 2.7 mM KCl, 0.5 mM MgCl_2_). Where applicable, cells were treated with inhibitors (Concanamycin A at 100 ng/mL (Santa Cruz CAS 80890-47-7) or Cytochlasin D at 10 µg/mL (ThermoFisher NC9996786). Approximately 2-5 beads per macrophage were added and fluorescence was measured using a Synergy Biotek plate reader set at 37°C with 5% CO_2_. Fluorescence for carboxyfluorescein beads was measured every 2 minutes for 2 hours at excitation 450 nm/emission 520 nm and excitation 490 nm/emission 520 nm. Fluorescence for DQ-BSA/AF594 beads was measured every 2 minutes for 4 hours at excitation 490 nm/emission 520 nm (DQ-BSA) and excitation 590 nm/emission 617 nm (AF594).

### Macrophage Stimulation and RNA-sequencing

Differentiated, lifted macrophages were plated into 6-well plates at 1.0 x 10^6^ cells per well (5.0 x 10^5^ cells/mL) and allowed to adhere overnight at 37°C, 5% CO_2_. The following day the cells were stimulated with *B. burgdorferi* B31, MOI 100 or 100 ng/mL *S. enterica* serovar Minnesota LPS (Invivogen tlrl-smlps) for 10 hours at 37°C, 5% CO_2_. Cells were scraped, washed, and preserved in RNAlater (ThermoFisher AM7020). Samples were incubated in RNAlater overnight at 4°C before storage at -80°C. RNA was extracted using the miRNAeasy isolation kit with Qiazol (Qiagen). DNA was depleted using TURBO DNase (Invitrogen AM2238) and RNA repurified using the Monarch® Spin RNA Cleanup Kit (New England Biolabs T2050S). Samples were submitted to Azenta Life Sciences/Genewiz for RNA quality assessment, library preparation, and sequencing (Illumina NovaSeq X+, 2×150 bp). Ribosomal RNA were depleted by polyA enrichment. We obtained an average of 21,607,624 paired-end reads per sample (range: 17,840,624 to 26,411,341 paired-end reads).

### RNA-sequencing Analysis

All analyses were performed separately for *P. leucopus* and *M. musculus*. Published *P. leucopus* (GCF_004664715.2) [23] and *M. musculus* (GRCm39) genomes were used for read mapping using STAR version 2.6.1 [40]. Gene expression was summarized using RSEM version 1.3.1 [41]. Annotations were based on the NCBI annotated genes. All differential gene expression analyses were performed with R (version 4.3.1) using DESeq2 version 1.40.2 [42] using standard parameters. Only genes with ≥10 reads across all conditions (summed) were included in the analysis. Differential gene expression, modeling gene ∼ condition, was determined using DESeq2, where the condition term differentiated macrophages based on mock treatment, treatment with LPS, or *B. burgdorferi* infection. Corrected p-values were generated using a Wald test and a Nejamini-Hochberg calculated false discovery rate. Genes with an adjusted p-value ≤ 0.05 were considered significant. All differential gene expression data is included in **Supplemental Files 2-5**. Pathway analysis was performed with the use of QIAGEN Ingenuity Pathway Analysis (QIAGEN Inc., https://digitalinsights.qiagen.com/IPA) [43] using genes with an adjusted p-value cutoff of <0.05.

### Data Availability

RNA sequencing data are available on the NCBI Gene Expression Omnibus (GEO Accession: GSE283617). All cells, plasmids, and reagents in this study are freely available upon request. For lentivirus constructs please contact Dr. Stephen C. Bunnell.

## RESULTS

### Development of a *P. leucopus* BMDM Differentiation Method

For many years innate immune research has relied on recombinant or cell line-derived M-CSF to differentiate monocyte cell populations from mouse and human samples [44, 45]. However, these tools have not yet been validated and/or generated for use with *P. leucopus*. Amino acid sequence identity between *M. musculus* and *P. leucopus* M-CSF is quite high (81.9% Identity) **(Figure 1A),** indicating that it could be possible to differentiate *P. leucopus* BMDMs in a culture media containing *M. musculus* M-CSF from L-929 supernatant as is frequently performed for *M. musculus* BMDM generation [44, 46]. Firstly, differentiation of *P. leucopus* BMDM was attempted by utilizing murine M-CSF enriched L-929 cell supernatant on flushed bone marrow from C57BL/6J *M. musculus* and *P. leucopus*. C57BL/6J bone marrow examined by phase contrast microscopy over a week of differentiation demonstrated changes in morphology and an increase in density consistent with BMDM differentiation, however the same was not observed with bone marrow from *P. leucopus* **(Figure 1B)**. Thus, we conclude that *M. musculus* M-CSF is insufficient to differentiate *P. leucopus* bone marrow into BMDMs. We speculated that a *P. leucopus*-specific M-CSF could have higher differentiation efficiency. To generate *P. leucopus* M-CSF, we transduced a human codon optimized, pCMV-driven *P. leucopus Csf1* gene into HEK293T cells via self-inactivating lentivirus [30] **(Figure 1C)**. *P. leucopus* bone marrow was incubated with media enriched with supernatant from *P. leucopus* M-CSF-expressing HEK293T cells (HEK293T^M-CSF^) over six days. Observation by phase contrast microscopy showed similar changes in cell proliferation, morphology, and adhesion to the plate as C57BL/6J bone marrow incubated with L-929 enriched media **(Figure 1B, 1D)**. To compare the relative changes in cell size and quantity during differentiation, BMDMs produced by either enriched media from L-929 cells or HEK293T^M-CSF^ were measured by flow cytometry at a fixed sample volume. There were species specific differences in the extent of cell proliferation, but both enriched media produced analogous populations of similar sizes by forward scatter histogram **(Figure 1E)**. To confirm differentiated cell identity, we performed reverse transcription quantitative PCR (RT-qPCR) to assess expression of hematopoietic progenitor cell (HPC) and monocyte markers—*Kit* and *Cd68,* accordingly. L-929 supernatant was able to induce *Cd68* in C57BL/6J bone marrow-derived cells **(Figure 1F)** and HEK293T^M-CSF^ supernatant was able to induce expression in *P. leucopus* bone marrow-derived cells **(Figure 1G).** Similarly, differentiating C57BL/6J cells had reduced expression in *Kit* **(Figure 1H),** and a similar trend was observed with *P. leucopus* (p=0.09, **Figure 1I).** Together, these data suggest that our HEK293T^M-CSF^ cells are capable of inducing *P. leucopus* BMDMs, comparable to L-929-driven C57BL/6J BMDM generation.

**Figure 1:**
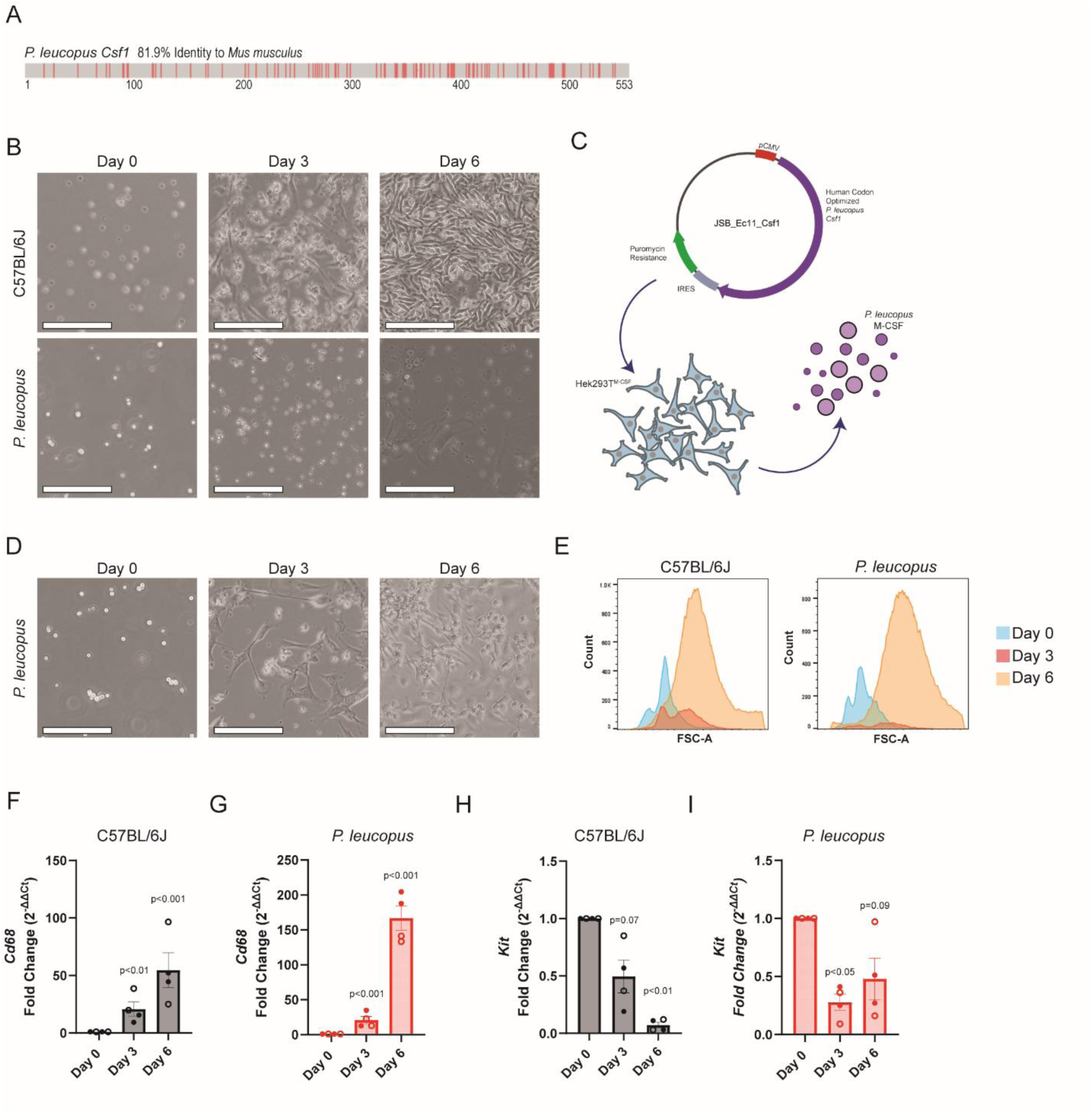
*P. leucopus*-specific M-CSF enables differentiation of bone marrow into macrophages. (A) *P. leucopus Csf1* shares substantial amino acid identity (81.9%) with *M. musculus* M-CSF. Differences in amino acid sequence are depicted with red dashes. Gaps are depicted by white space. Aligned amino acids are depicted in gray. (B) L-929 supernatant is able to induce C57BL6/6J bone marrow differentiation into macrophages, but not *P. leucopus* bone marrow. Phase contrast microscopy was taken at days 0, 3, and 6 of the differentiation procedure. (C) Schematic of *P. leucopus* M-CSF generation. Generated using Biorender.com and the NIH BioArt Source. (D) Supernatant from HEK293T^M-CSF^ cells is able to induce *P. leucopus* bone marrow differentiation into macrophages. (E) C57BL/6J bone marrow cells treated with L-929 supernatant or *P. leucopus* bone marrow cells treated with HEK293T^M-CSF^ supernatant undergo changes in cell abundance and size. Flow cytometry was performed on a fixed volume of cells at days 0, 3, and 6 of the differentiation procedure to measure the FSC-A. (F-I) C57BL/6J bone marrow cells treated with L-929 supernatant or *P. leucopus* bone marrow cells treated with HEK293T^M-CSF^ supernatant upregulate *Cd68* (F, G) and downregulated *Kit* (H,I). RT-qPCR was performed six days into the differentiation procedure. p-values were calculated by a one sample t-test comparing the log transformed fold change values to a predicted value of 0. Panels B, D, and E are representative of four independent experiments. Panels F-I summarize four experiments, where each point represents an individual mouse. Closed circles represent males and open circles represent females. Bars mark the mean, error bars the standard error of the mean. Phase contrast micrographs were captured using an ECHO Rebel. For Panels B and D, the scale bar represents 135µm.

### *P. leucopus* BMDMs have similar polarization under unstimulated conditions to C57BL/6J BMDMs

In order to understand how *P. leucopus* BMDMs compare to C57BL/6J BMDMs in their behavior and response to pathogenic stimuli, we first wanted to ensure that technical differences in the differentiation protocols did not bias the macrophages towards different inflammatory phenotypes at baseline. To examine this question, we used RNA sequencing to compare the transcriptomes of *P. leucopus* BMDMs (n=3) and C57BL/6J BMDMs generated with L-929 supernatant (n=3) under unstimulated conditions **(Supplemental File 6, Supplemental File 7)**. We found that principal component analysis could distinguish between the two species based on their transcriptome by the first principal component (56.52% of the variation), indicating that transcriptional differences across the cells exist **(Figure 2A)**. However, we observed no statistically significant differences in M1-like and M2-like polarization markers [47, 48] **(Figure 2B)**, leading us to conclude that the differentiation protocols generate macrophages that are sufficiently similar to enable direct comparison. While M1 and M2 macrophage classifications simplify the heterogeneity of potential macrophage responses, the use of these identifiers provides a useful heuristic for determining species-specific differences between BMDMs derived *ex vivo* from a naïve cell population [49, 50].

**Figure 2:**
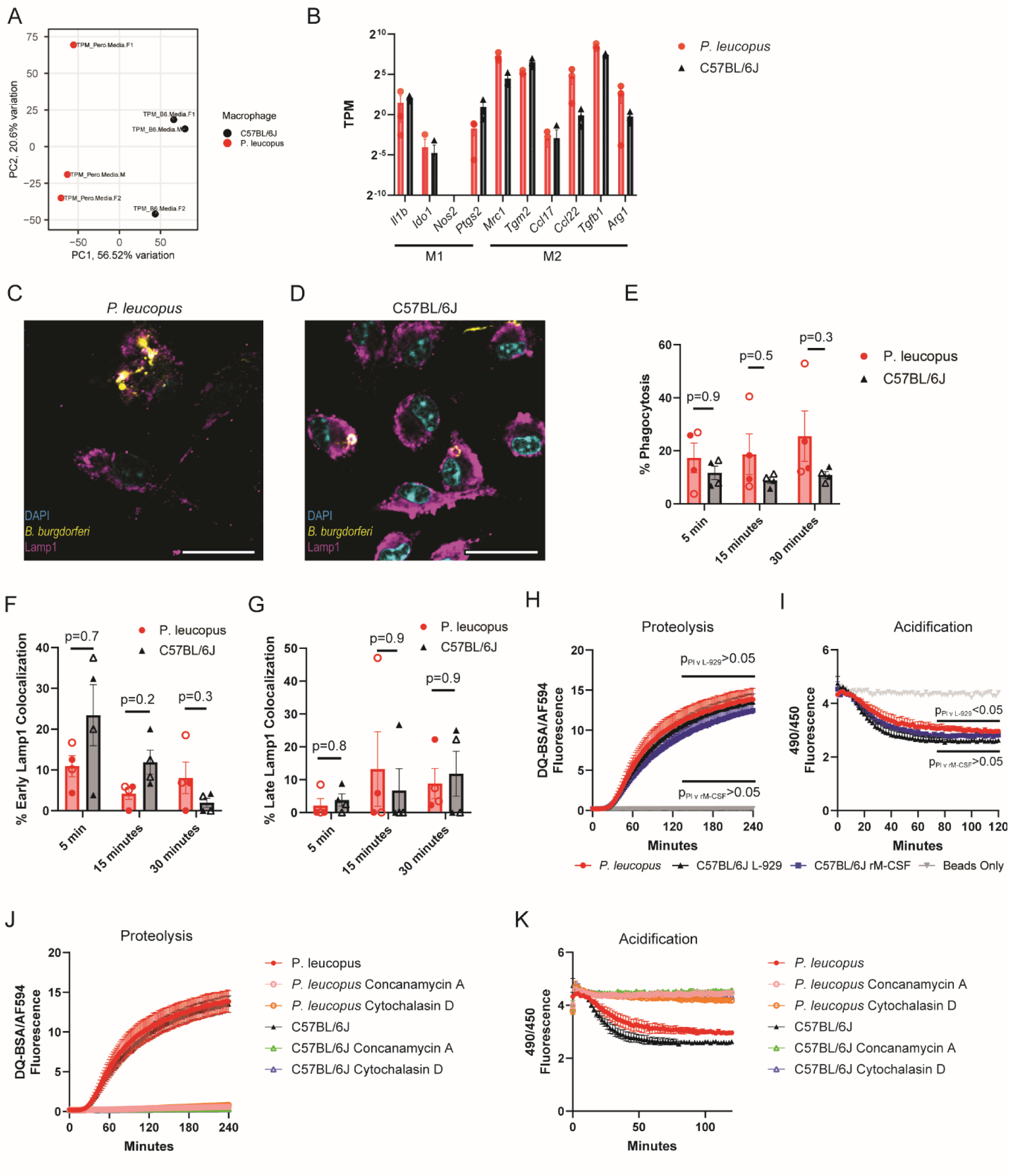
*P. leucopus* and C57BL/6J BMDMs display similar behaviors during baseline conditions. (A) Principal component analysis can distinguish between *P. leucopus* and C57BL/6J BMDM transcriptomes at baseline. RNA sequencing performed on three sets of *P. leucopus* and C57BL/6J BMDMs and analyzed by principal component analysis. (B) M1 and M2 polarization markers [47, 48] are similar between the two species at baseline. Each dot represents the transcripts per million for each gene under baseline conditions. No statistically significant differences in expression were observed by performing multiple t tests on the log transformed values with a Holm-Šídák test to correct for multiple comparisons. (C, D) *P. leucopus* and C57BL/6J BMDMs phagocytosis of *B. burgdorferi* and the colocalization with LAMP1 phagosome. Representative images taken 30 minutes post *B. burgdorferi* infection. Scale bar represents 22µm. (E-F) Quantification of microscopy does not reveal differences in *B. burgdorferi* phagocytosis or phagosomal maturation across species. Each point represents a single mouse, closed circles represent males and open circles represent females. Bars mark the mean, error bars the standard error of the mean. p-values calculated by 2-way ANOVA with Dunnett correction for multiple comparisons. (H-K) Analysis of *P. leucopus* proteolysis (H, J) and phagosome acidification (I, K). Fc receptor mediated phagocytosis of proteolysis and pH sensitive beads brings beads to the phagolysosome where we observe increased ratiometric signal from DQ-BSA proteolysis (H, J) and reduced ratiometric signal from acidic fluorochrome degradation (I, K). Proteolysis and acidification were measured for C57BL/6J using L-929 supernatant and recombinant M-CSF (rM-CSF). The ratio of the carboxyfluorescein fluorescence signal at excitation 490 nm (pH-sensitive) versus 450 nm (pH-insensitive) provides a readout of relative pH (I, K). Panels H and I include a “Beads Only” control where fluorescence was measured in wells containing beads but no macrophages. Inhibitor treatments of concanamycin A (100 ng/mL) prevent proteolysis (J) and acidification (K) in both species while cytochalasin D (10ug/ml) inhibits phagocytosis of the beads (J, K). For Panels J and K, C57BL/6J macrophages were differentiated with L-929 supernatant. p-values for H and I were generated by two-way ANOVA with Dunnett’s multiple comparison test. H-K represent summarized data from four experiments where symbols mark the mean and error bars the standard error of the mean.

### *P. leucopus* BMDMs have similar proteolysis but potentially altered phagosome acidification to C57BL/6J BMDMs

The ability for macrophages to phagocytose and process pathogens is critical to the innate immune system’s ability to restrain infection. Not only do these processes result in pathogen destruction, but phagosomal maturation and proteolysis influence subsequent immune responses reliant on ligand detection by pattern recognition receptors and antigen processing for display by the major histocompatibility complex [51, 52]. We sought to examine the ability of *P. leucopus* macrophages to phagocytose *B. burgdorferi* and the rate at which the phagosome becomes associated with LAMP1, which marks stages of lysosomal development [33, 36]. Phagocytosis was examined by confocal microscopy after infecting BMDMs with *B. burgdorferi* for 5, 15, and 30 minutes **(Figure 2C**, **2D)**. No significant differences in total phagocytosis or LAMP1 colocalization were observed between species **(Figure 2E-2G)**.

We further examined phagosomal maturation and activity in the species. We utilized a pair of silica bead-based systems with pH-sensitive carboxyfluorescein and dye-quenched bovine serum albumin (DQ-BSA)/AF594 conjugates to assess acidification and proteolysis respectively [39]. Co-incubation of the DQ-BSA/AF594 beads with BMDMs demonstrated a lack of statistically different proteolysis between the two species **(Figure 2H).** Despite *P. leucopus* and C57BL/6J proteolytic similarities, incubation of the pH sensitive carboxyfluorescein beads with *P.* leucopus BMDMs showed slightly less phagosomal acidification, indicating that phagosomal maturation may be different than in C57BL/6J BMDMs **(Figure 2I)**. We also performed both assays with C57BL6/J BMDMs generated with recombinant M-CSF and did not find a statistically significant difference between these cells and *P. leucopus* BMDMs for either phenotype (**Figure 2H, I)**, making it unclear whether this subtle acidification phenotype plays a role in driving interspecies differences. We found these processes were sensitive to concanamycin A (100 ng/mL) and cytochalasin D (10 µg/mL) in both species **(Figure 2J, K).**

### *P. leucopus* and C57BL/6J BMDMs exhibit modest differences in their transcriptomic profiles following *B. burgdorferi* infection

We next sought to understand whether we could observe differences in the immunological response of *P. leucopus* and *M. musculus* BMDMs to bacterial stimuli. To do this, we performed a small pilot RNA sequencing experiment in which BMDMs from three rodents per species (two females, one male) were stimulated either with *B. burgdorferi* B31 or LPS for ten hours.

Following *B. burgdorferi* stimulation, principal component analysis clearly separated macrophages from *P. leucopus* or C57BL/6J mice along the first principal component (41.76% of the variation). Curiously, while the second component (22.59% of the variation) was able to separate uninfected and infected C57BL/6J macrophages, this separation was less successful for the *P. leucopus* BMDMs **(Figure 3A).** We quantified transcripts from 13,885 *P. leucopus* genes and found 3,174 differentially expressed genes (p_adj_≤0.05), including 1,662 upregulated genes and 1,512 downregulated genes **(Figure 3B, Supplemental File 2)**. We quantified transcripts from 15,282 *M. musculus* genes and found 5,614 differentially expressed genes – 2,856 upregulated and 2758 downregulated **(Figure 3C, Supplemental File 3)**. Restricting the analyses to genes quantified in both species allowed for examination of 9,806 genes – 2,521 of which were differentially expressed in *P. leucopus* BMDMs (1,342 upregulated, 1,179 downregulated) and 4,171 differentially expressed in C57BL/6J BMDMs (2,163 upregulated, 2,008 downregulated) **(Supplemental File 8)**. Examining M1-like and M2-like polarization markers [47, 48] revealed a modest shift towards M1-like polarization among *P. leucopus* and C57BL/6J macrophages following *B. burgdorferi* infection **(Figure 3D)**. Using the QIAGEN Ingenuity Pathway Analysis tool [43] revealed robust evidence of numerous canonical pathways becoming activated in both species, including interleukin-1 family signaling and NF-κB signaling, but few pathways that exhibited differences in activation across the species **(Figure 3E, Supplemental File 9)**.

**Figure 3:**
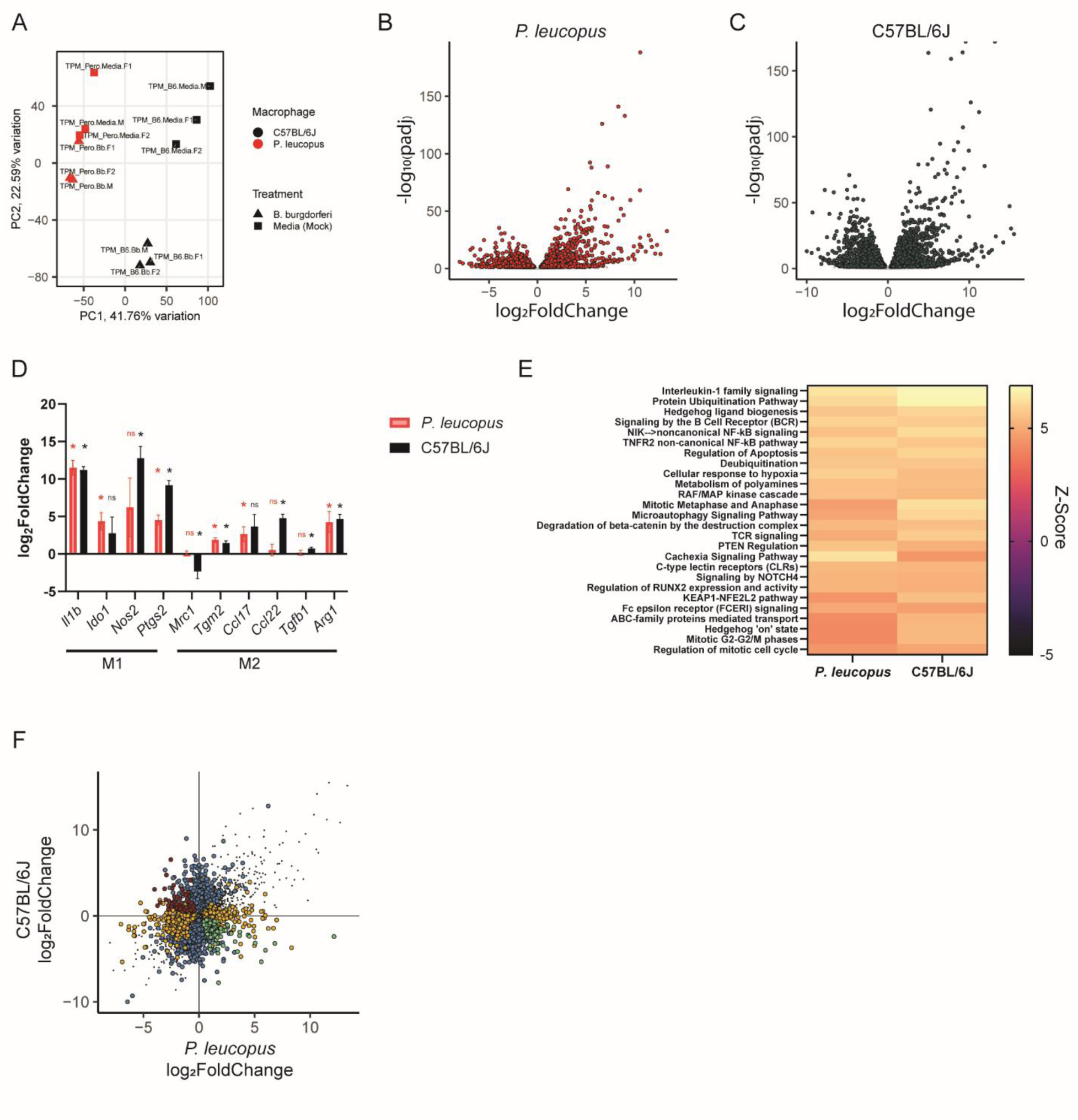
Comparison of *P. leucopus* and C57BL/6J BMDM responses to *B. burgdorferi* B31. (A) Principal component analysis is able to distinguish between *P. leucopus* and C57BL/6J BMDMs following *B. burgdorferi* infection (strain B31, multiplicity of infection 100, ten-hour stimulation). (B, C) Multiple genes are differentially expressed in (B) *P. leucopus* or (C) C57BL/6J BMDMs following *B. burgdorferi* stimulation. Adjusted p values (padj) are calculated based on a false discovery rate. Genes with an adjusted p value ≤ 0.05 are depicted as red or dark gray dots. (D) *P. leucopus* and C57BL6/J BMDMs display similar changes in expression of M1 and M2 polarization markers [47, 48] following *B. burgdorferi* infection. Listed p-values are the same FDR corrected p-values generated in panels B and C. (E) *P. leucopus* and C57BL/6J display activation of similar pathways following *B. burgdorferi* infection. Pathway analysis was performed with the QIAGEN Ingenuity Pathway Analysis tool [43] using a p-value cutoff of <0.05. Top 25 pathways are depicted here. All plotted values have Z-scores > 2, suggesting activation of the pathways. (F) *P. leucopus* and C57BL/6J BMDMs display some differences in gene expression following *B. burgdorferi* infection. All genes with a padj≤0.05 in one or both conditions are plotted according to their log_2_ fold change in each species Dots in gray are differentially expressed in the same direction in each species. Dots in blue are differentially expressed in C57BL/6J BMDMs but not differentially expressed (padj>0.05) in *P. leucopus* BMDMs. Dots in orange are differentially expressed in *P. leucopus* BMDMs but not differentially expressed (padj>0.05) in *P. leucopus* BMDMs. Dots in red and green are differentially expressed in opposite directions across the two species. All data are from three independent experiments with three rodents (2 females, 1 male) from each species.

Based on p-value cutoffs (p_adj_≤0.05), numerous genes were differentially expressed in one species but either (a) not differentially expressed or (b) exhibited an opposite direction of effect in the other species (790 genes were uniquely upregulated in C57BL/6J BMDMs, 304 genes were uniquely upregulated in *P. leucopus* BMDMs, 986 genes were uniquely downregulated in C57BL/6J BMDMs, and 455 genes were uniquely downregulated in BMDMs) **(Figure 3F, Supplemental File 8)**. During manual examination of genes upregulated specifically in *P. leucopus* BMDMs we noted multiple genes involved in inflammation, including *Ccl17, Il1r1*, *Il1r2*, *Il1rl1*, *Il18*, *Gata3*, *Ncf1*, *Tifa*, *Ifnar1*. Manual examination of immunity-associated genes upregulated in *M. musculus* but not *P. leucopus* BMDMs include *Ccl22*, *Il17ra*, *Il17rd*, *Il20rb*, *Il2rg*, *Il7*, *Mcoln2*, *Traf2*, *Traf5*, *Trim13*. We note that examining the 77 genes specifically with differential expression in opposite direction of effect across the two species revealed only a few additional genes canonically associated with immunity, including *Casp1* (upregulated in C57BL/6J, downregulated in *P. leucopus*), *Sod3* (upregulated in C57BL/6J, downregulated in *P. leucopus*), *Il15* (upregulated in *P. leucopus*, downregulated in C57BL/6J), and *Casp3* (upregulated in *P. leucopus*, downregulated in C57BL/6J).

### *P. leucopus* and C57BL/6J BMDMs exhibit modest differences in their transcriptomic profiles following LPS stimulation

Comparing *P. leucopus* and C57BL/6J BMDMs stimulated with LPS revealed many of the same observations as comparing the species following *B. burgdorferi* stimulation. Principal component analysis was able to separate the species along the first principal component (40.96% of the variation), but was only able to distinguish between stimulation status for C57BL/6J macrophages along the second principal component (19.68% of the variation) **(Figure 4A)**. Transcripts for 13,599 *P. leucopus* genes were quantified, and 1,734 genes were upregulated in response to LPS while 1,493 genes were downregulated **(Figure 4B, Supplemental File 4)**. Of 14,967 genes quantified in C57BL/6J macrophages, 2,530 were upregulated in response to LPS and 2,094 genes were downregulated **(Figure 4C, Supplemental File 5)**. Restricting the analyses to genes that were differentially expressed in both datasets limited the analysis to 9,691 genes – 2,436 were differentially expressed in *P. leucopus* BMDMs (1,308 upregulated, 1,128 downregulated) and 3,379 were differentially expressed in C57BL/6J BMDMs (1,805 upregulated, 1,574 downregulated) **(Supplemental File 10)**. Both *P. leucopus* and C57BL/6J BMDMs displayed a modest shift towards M1-like associated gene expression following stimulation (**Figure 4D)**. Leveraging the QIAGEN Ingenuity Pathway Analysis tool [43] again revealed strong evidence of Interleukin-1 family signaling and NF-κB signaling, as well as activation of other immune-related pathways, but very few differences between the two species **(Figure 4E, Supplemental File 9)**.

**Figure 4:**
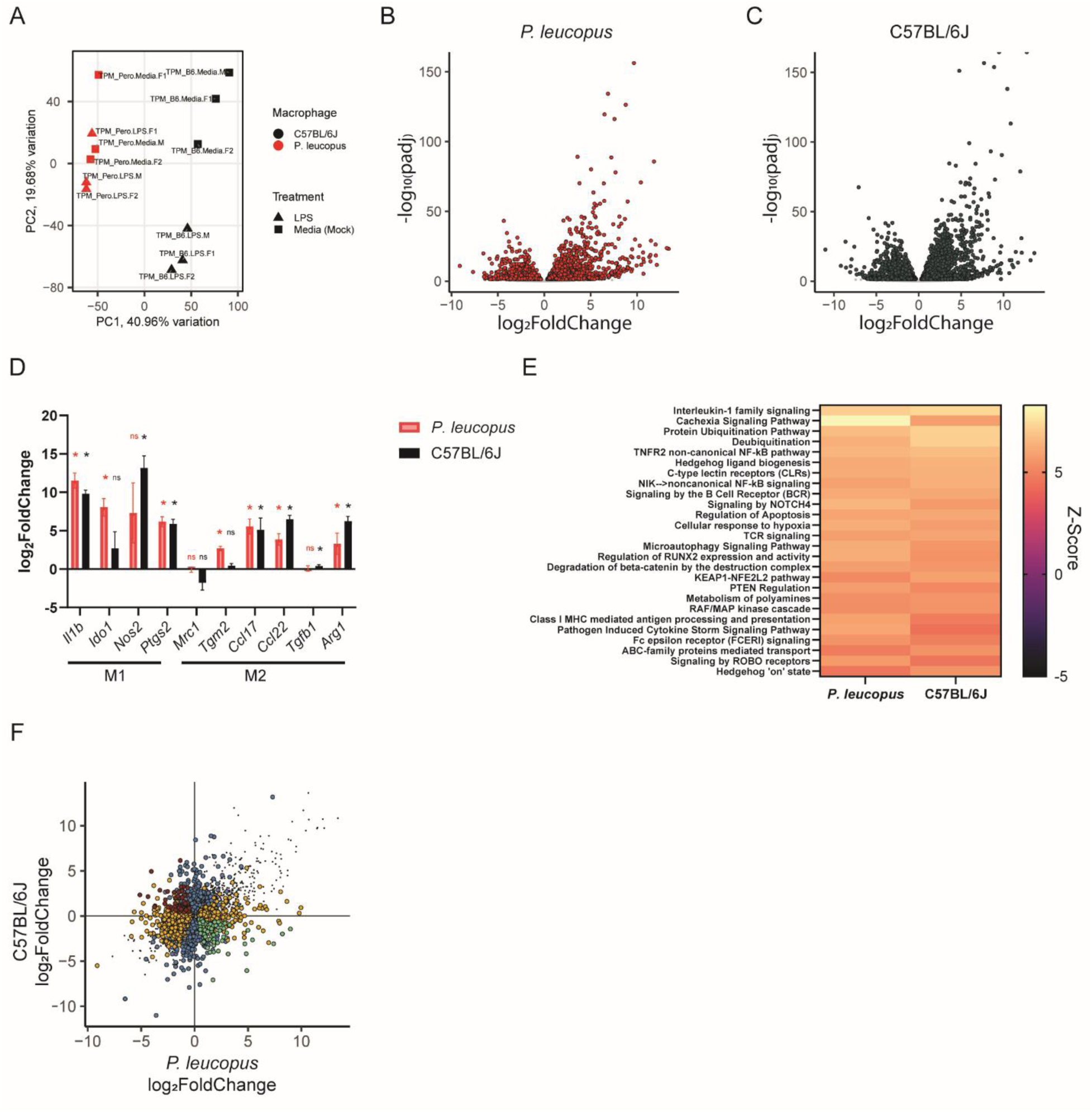
Comparison of *P. leucopus* and C57BL/6J BMDM responses to LPS. (A) Principal component analysis is able to distinguish between *P. leucopus* and C57BL/6J BMDMs following LPS stimulation (ten-hour stimulation). (B, C) Multiple genes are differentially expressed in (B) *P. leucopus* or (C) C57BL/6J BMDMs following LPS stimulation. Adjusted p-values (padj) are calculated based on a false discovery rate. Genes with an adjusted p-value ≤ 0.05 are depicted in red or dark gray. (D) *P. leucopus* and C57BL6/J BMDMs display similar changes in expression of M1 and M2 polarization markers [47, 48] following LPS stimulation. P-values are the same FDR corrected p-values in panels B and C. (E) *P. leucopus* and C57BL/6J display activation of similar pathways following LPS stimulation. Pathway analysis was performed with the QIAGEN Ingenuity Pathway Analysis tool [43] using a p-value cutoff of <0.05. The top 25 pathways are depicted here. All plotted values have Z-scores > 2, suggesting activation of the pathways. (F) *P. leucopus* and C57BL/6J BMDMs display some differences in gene expression following LPS stimulation. All genes with a padj≤0.05 in one or both conditions are plotted according to their log_2_ fold change in each species. Dots in gray are differentially expressed in the same direction in each species. Dots in blue are differentially expressed in C57BL/6J BMDMs but not differentially expressed (padj>0.05) in *P. leucopus* BMDMs. Dots in orange are differentially expressed in *P. leucopus* BMDMs but not differentially expressed (padj>0.05) in *P. leucopus* BMDMs. Dots in red and green are differentially expressed in opposite directions across the two species. All data are from three independent experiments with three rodents (2 females, 1 male) from each species.

Similar to our analysis of *B. burgdorferi*-induced gene expression changes, we next examined genes that were differentially expressed in one species but not the other and/or exhibited opposite direction of differential expression in response to LPS **(Figure 4F, Supplemental File 10)**. We identified 611 genes that were uniquely upregulated and 709 genes that were uniquely downregulated in *P. leucopus* BMDMs, as well as 1,108 genes that were uniquely upregulated and 1,155 genes uniquely downregulated in C57BL/6J BMDMs. We note multiple interesting trends, including that *Casp3*, *Il10, Il10ra*, *Il15*, *Il17ra*, *Il1r2*, *Il1rl1*, *Il23a*, *Rab11a*, *Rab8b*, *Tlr7,* and *Traf7* were upregulated in *P. leucopus* but not C57BL/6J. Conversely, *Casp1*, *Casp7*, *Casp8*, *Il15ra*, *Il17rd*, *Il20rb*, *Il21r*, *Il27*, *Il2rg*, *Il36a*, *Il7*, *Jak3*, *Syk*, and *Tlr3* were upregulated in C57BL/6J, but not in *P. leucopus*. 98 genes were upregulated in *P. leucopus* but downregulated in C57BL/6J BMDMs and 84 genes were upregulated in C57BL/6J but downregulated in *P. leucopus*.

### Comparison of BMDM gene expression in response to LPS with rodent and fibroblast studies

Substantial past work has sought to understand *P. leucopus* transcriptomic responses to LPS, and so we sought to examine how well our *ex vivo* phenotypes could recapitulate past findings. Specifically, we compared our data from *P. leucopus* and *M. musculus* BMDMs to a previous study by Milovic *et al*. of *P. leucopus* and *M. musculus* RNA sequencing of blood following LPS injection (four hours post-injection) [10]. Restricting the analysis to genes that were differentially expressed both in our dataset and the Milovic *et al.* dataset revealed a positive association (r^2^=0.19) with *P. leucopus* **(Figure 5A)** and a weak but still statistically significant relationship with outbred CD-1 *M. musculus* (r^2^=0.029) **(Figure 5B)**. We next examined the association between the LPS-induced transcriptome in *P. leucopus* BMDMs (ten-hour stimulation, **Figure 4**) and previous results obtained by Balderrama-Gutierrez *et al*. with skin fibroblasts (twenty-four hour stimulation, [11]). Examining genes that were differentially expressed in both datasets revealed a weak (r^2^=0.038) association between BMDM and skin fibroblast gene expression **(Figure 5C)**. Of note, the fibroblast dataset has a weaker association (r^2^=0.026) **(Figure 5D)** with differential gene expression in the Milovic *et al. P. leucopus* dataset than we observed with BMDMs (r^2^=0.19) **(Figure 5A)**. This appears to be driven by: (1) a variable number of differentially expressed genes shared between the datasets (1,186 between macrophages and blood compared to 238 genes between fibroblasts and blood), and (2) substantially more volatile gene expression changes in fibroblasts following stimulation (for example, *Nfkb2* has a log_2_-fold change of 2.35 in blood, compared to 2,569.39 in fibroblasts). While these differences could be attributed to differences in exposure time to LPS, they may also highlight differences in LPS processing and responsiveness between non-immune cell subtypes (fibroblasts) and immune cells (peripheral blood cells or BMDMs).

**Figure 5:**
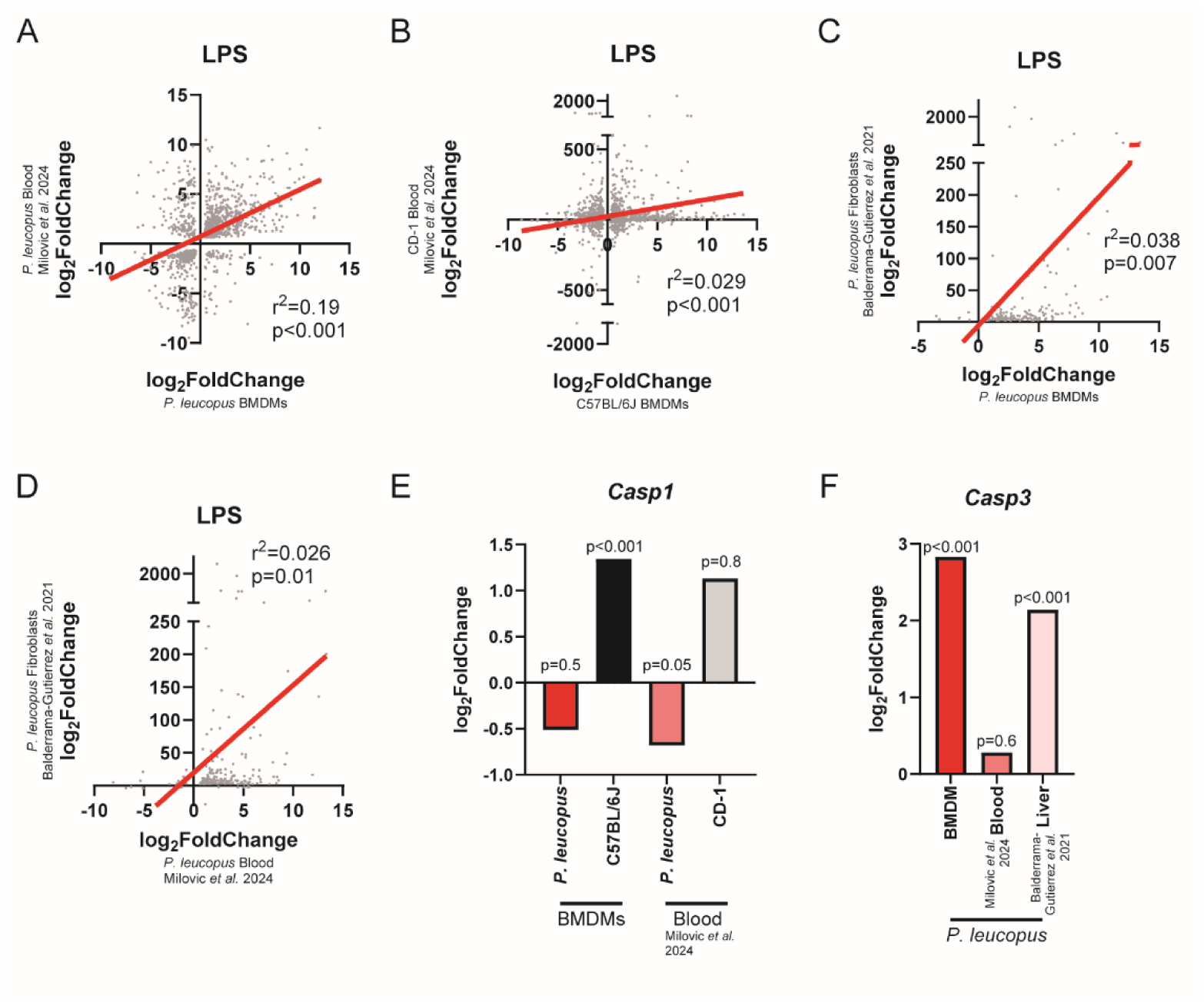
Comparison of *P. leucopus* BMDM responses to LPS with previously reported *in vivo* phenotypes. (A-B) There is a positive correlation between gene expression changes observed with LPS stimulation in (A) *P. leucopus* and (B) C57BL/6J BMDMs and the blood transcriptomes following LPS injection [10]. (C) There is a weak positive correlation between *P. leucopus* BMDM and fibroblast gene expression changes [11] following LPS stimulation. (D) There is a weak positive correlation between *P. leucopus* fibroblast and *P. leucopus* blood gene expression [10] following LPS stimulation. For panels A-D only genes differentially expressed in both datasets were examined. r^2^ and p-values were calculated based on a simple linear regression (plotted in red). (E) *Casp1* gene expression is higher in C57BL/6J macrophages and lower in *P. leucopus* blood following LPS stimulation [10]. (F) *Casp3* gene expression is elevated in *P. leucopus* BMDMs and livers [11], but not blood [10] following LPS stimulation. For panels E and F, p-values for BMDM gene expression were generated by the FDR corrected p-values. p-values for blood and liver are based on the reported p-values from the publications from which they were derived [10, 11].

Of particular note, in the above section we found that *Casp1* expression was upregulated in C57BL/6J macrophages, but not differentially expressed in *P. leucopus* macrophages following LPS stimulation **(Figure 5E)**. Interestingly, *P. leucopus* appear to downregulate *Casp1* in blood following LPS injection, while there is no difference in expression in CD-1 *M. musculus* [10] **(Figure 5E)**. Thus, the relative relationship between *Casp1* expression across the species is preserved in BMDMs and blood. Curiously, another component of the pyroptosis pathway, *Nlrp3*, is upregulated in both C57BL/6J and *P. leucopus* macrophages, but others have found that CD-1 mice downregulate the gene while *P. leucopus* upregulate it in blood during LPS exposure [10]. Relatedly, *Casp3* was found to be upregulated in *P. leucopus*, but unchanged in C57BL/6J BMDMs **(Figure 5F)**. However, in animals, while *Casp3* was not differentially expressed in *P. leucopus* blood in response to LPS [10]—other work found it was specifically upregulated in *P. leucopus*, and not BALB/c *M. musculus*, livers during LPS treatment (Four hours post injection) [11] **(Figure 5F)**. Taken together, these data suggest that there may be cross-species differences in pyroptosis pathway activation during LPS stimulation.

## DISCUSSION

To our knowledge, this work represents the first published use of bone marrow-derived macrophages for any *Peromyscus* species. Numerous zoonotic pathogens (*B. burgdorferi*, *Borrelia miyamotoi, Borrelia bissettii*, Powassan Virus, *Anaplasma phagocytophilum*, *Babesia microti*, *Ehrlichia muris*, *Bartonella* species, and Hantavirus, among others) are confirmed or suspected to be present in *P. leucopus* as a reservoir [2–8], meaning that the host-pathogen interactions in this species have direct impacts on human health by enabling zoonotic spillover. Therefore, expanding the toolkit to study immunological responses in *P. leucopus* represents an opportunity to better understand how pathogens colonize and transmit through this rodent.

In this work, we benchmarked *P. leucopus* BMDM behavior to the well-studied C57BL/6J *M. musculus* strain. We determined that M-CSF secreted by HEK293T^M-CSF^ was sufficient to differentiate BMDMs from *P. leucopus* in a manner similar to L-929 conditioned media used on *M. musculus* BMDMs. We qualified this by changes in cell size, expression of genes related to cell markers *Cd68* and *Kit*, and visual comparisons by phase contrast microscopy. This novel HEK293T^M-CSF^ cell pool will be instrumental for future differentiation of *P. leucopus* monocytes for study of *Borrelial* infection as well as other zoonotic pathogens that canonically infect *P. leucopus*. While the application of this method is in its infancy, these results are an important step for utilizing *P. leucopus* as a model organism in a research setting.

Stimulation of BMDMs with *B. burgdorferi* and LPS demonstrated interesting transcriptional changes both unique and shared between stimulations and species. Many genes upregulated by *P. leucopus* during *B. burgdorferi* stimulation are related to Il-1β receptors, and other inflammation related receptors and transcription factors like *Ifnar1* and *Gata3*. Comparatively, C57BL/6J responses to *Bb* included upregulation of pyroptosis-related genes such as *Casp1*and *Il1b*, as well as multiple cytokines and chemoattractants typically associated with T-helper cell activity, and specifically Th17 cells. Interestingly *P. leucopus* downregulates *Casp1* in response to *B. burgdorferi* while still upregulating *Il1b*, indicating that *P. leucopus* are potentially translating enough pro-Il-1β to reliably carry out pyroptosis, but downregulating the transcription of *Casp1* to reduce the amount of Caspase-1, preventing the cleavage reaction that creates immunologically active Il-1β [53, 54].

Under LPS stimulation many proinflammatory pathways are induced between both *P. leucopus* and C57BL/6J, with some notable upregulations of *Rab11a* and *Rab8b* in *P. leucopus*, showing enhanced expression of chaperones associated with endosomal recycling and exocytosis [55, 56] as well as involvement with endosomal trafficking and immune signaling upon phagocytosis of *B. burgdorferi* [33]. Additionally, *P. leucopus* upregulates expression of *Il10* and *Il1ra* in response to LPS many times more than C57BL/6J upregulation of *Il10*. C57BL/6J transcriptional changes also included the upregulation of *Casp1*, *Casp7*, *Casp8*, various Il2 family cytokines and receptors, and kinases required for further activation of receptor signaling domains such as *Syk* and *Jak3*.

We note that our intent is for this work to serve as a launching point for future studies, rather than a comprehensive examination of *P. leucopus* innate immunity. As such, we are aware of numerous limitations of the work presented here. First, we relied on transcriptomic data as a proxy for immune signaling as there are limited commercially available antibodies that recognize *P. leucopus* proteins, meaning ELISA, western blotting, and flow cytometry are difficult in these systems. Further, our work here used members from a single colony of *P. leucopus* and a single strain of *M. musculus*, meaning that the results presented here may not be representative of their species as a whole. During our RNA sequencing experiments, we used single doses and timepoints to interrogate the immune response to bacterial stimuli. Future work may consider a more complete dose-response or time course that would provide another set of dimensions for interpretation. Further, *P. leucopus* and C57BL/6J macrophages both upregulate *Acod1*, encoding aconitate decarboxylase, in response to *B. burgdorferi* or LPS. Past work has demonstrated that this gene is a negative regulator of inflammatory processes and contributes to the induction of innate immune tolerance [57–59]. Examining whether innate immune tolerance occurs in *P. leucopus* and how this compares to C57BL/6J and human tolerance could be interesting for future studies.

Bacterial strain is another important consideration for immune stimulation. The *B. burgdorferi* B31 wild-type strain is a common choice for performing bacterial stimulations, however, it may not be representative of the *B. burgdorferi* strains that have co-evolved with *P. leucopus*. Recent work has demonstrated that different *B. burgdorferi* lineages are more or less likely to successfully use *P. leucopus* as a reservoir [16, 26, 60], which correlates with *in vitro* phenotypes [26]. Thus, future work may benefit from screening divergent responses of these macrophages to different *B. burgdorferi* strains. Beyond *B. burgdorferi*, other zoonotic pathogens (Powassan virus, *Anaplasma phagocytophilum*, *Ehrlichia* spp) utilize phagocytes during host colonization and pathogenesis [6, 61, 62], meaning these tools could be used to better understand their interactions with *P. leucopus*. In particular, leveraging intraspecies natural host diversity in Powassan virus-macrophage studies has previously yielded insight into key host-pathogen interactions [63], and we hope the ability to use inter-species diversity could provide further insight into Powassan virus resistance and pathogenesis. Finally, the M-CSF producing cells generated here could be used to differentiate monocytic cells from the periphery rather than common monocyte progenitors from the bone marrow. The stimuli experienced by cells after hematopoiesis is crucial for the diversity of macrophage responses that contribute to disease phenotypes, and while our investigation of naïve cells highlights implicit differences between species, even more may be understood by examining *P. leucopus* innate phenotypes with greater breadth.

In conclusion, we believe that this work will unlock numerous exciting studies in order to understand how zoonotic pathogens are tolerated by reservoir hosts in nature before spilling over into humans. We hope that the tools generated here will prove useful to others in the field studying innate immunity in this critical human-interacting animal.

## ACKNOWLEDGEMENTS

We thank Tufts Comparative Medical Services for maintenance of the rodents used in this study. Confocal Microscopy was performed using the Tufts Center for Neuroscience Imaging and Cell Analysis Core. Flow cytometry was performed with help from the Tufts School of Medicine Flow Cytometry Core. The plasmid schematic in Figure 1 was generated using graphics from Biorender.com and the NIH BioArt Source. The plasmid schematic in Supplemental File 1 was generated on Benchling.com. We thank Dr. Ryan Finethy for advice on bone-marrow derived macrophage cultivation, as well as past and present members of Dr. Linden Hu’s lab for feedback and support (particularly Dr. Peter Gwynne, Dr. Cecilia Silva, Kee-Lee Stocks, Yuelin Zhong, Stephanie You, Muskan Shrestha, Emily Vu, and Abigail Rivera Seda). Support from the National Institutes of Health to JSB (F32AI179104), LTH (R01AI150157, R01AI152210), and SRT (R01AI152209) enabled this project. TP is supported by the Global Lyme Alliance. Funders had no role in the design or interpretation of these studies.

## SUPPLEMENTAL MATERIAL LEGENDS

**Supplemental File 1**: Plasmid map of JSB_Ec11 *P. leucopus* M-CSF producing construct

**Supplemental File 2**: *P. leucopus* BMDM differentially expressed gene expression analysis in response to *B. burgdorferi*

**Supplemental File 3**: C57BL/6J BMDM differentially expressed gene expression analysis in response to *B. burgdorferi*

**Supplemental File 4**: *P. leucopus* BMDM differentially expressed gene expression analysis in response to LPS

**Supplemental File 5**: C57BL/6J BMDM differentially expressed gene expression analysis in response to LPS

**Supplemental File 6**: Transcript Per Million data for C57BL/6J BMDMs

**Supplemental File 7**: Transcript Per Million data for *P. leucopus* BMDMs

**Supplemental File 8**: BMDM differentially expressed gene analysis, restricted to genes present in *P. leucopus* and C57BL/6J datasets, in response to *B. burgdorferi*

**Supplemental File 9**: QIAGEN Ingenuity Pathway Analysis for *B. burgdorferi* and LPS stimulated cells

**Supplemental File 10**: BMDM differentially expressed gene analysis, restricted to genes present in *P. leucopus* and C57BL/6J datasets, in response to LPS

